# Neural Processing of Naturalistic Audiovisual Events in Space and Time

**DOI:** 10.1101/2024.05.14.594145

**Authors:** Yu Hu, Yalda Mohsenzadeh

## Abstract

Our brain seamlessly integrates distinct sensory information to form a coherent percept. However, when real-world audiovisual events are perceived, the specific brain regions and timings for processing different levels of information remain less investigated. To address that, we curated naturalistic videos and recorded fMRI and EEG data when participants viewed videos with accompanying sounds. Our findings reveal early asymmetrical cross-modal interaction, with acoustic information represented in both early visual and auditory regions, while visual information only identified in visual cortices. The visual and auditory features were processed with similar onset but different temporal dynamics. High-level categorical and semantic information emerged in multi-modal association areas later in time, indicating late cross-modal integration and its distinct role in converging conceptual information. Comparing neural representations to a two-branch deep neural network model highlighted the necessity of early fusion to build a biologically plausible model of audiovisual perception. With EEG-fMRI fusion, we provided a spatiotemporally resolved account of neural activity during the processing of naturalistic audiovisual stimuli.

## 1 Introduction

Most visual scenes are accompanied by sounds. Thus, when we perceive these scenes, not only does light enter the eyes, but also sound waves are captured by the ears. These two types of sensory information are processed through separate channels, yet our brain effectively integrates these distinct inputs to form a coherent percept^1,2^. However, the neural basis underlying the perception of real-world audiovisual events remains poorly understood.

Many studies have examined how the human brain perceives the external world using either vision or audition alone. It has been shown that each sensory system is linked to specific areas of the brain, and sensory information is processed through hierarchical stages along designated neural pathways^3–6^. In vision, different levels of visual information are gradually resolved through the ventral visual stream^7^, with V1 for low-level orientation^8–10^, V2-V4 for mid-level curvature^11–14^, and lateral and ventral occipitotemporal cortex (OTC) for high-level category^15–23^. The dorsal visual stream is mainly accounted for the role in visually guided action^24–26^, but also found involved in object perception^27–31^. Visual hierarchical processing can be accomplished rapidly within the first few hundred milliseconds^32–34^. How neural representations unfold over time and across brain regions has been well characterized^35–41^. In audition, object information can be extracted from sounds through hierarchical processing along the auditory pathway^42^. Acoustic features are first represented in the primary auditory cortex (A1) and later in the belt and para-belt regions, and finally high-level categorical information emerges in middle-to-posterior temporal cortex^5,43–47^. The processing speed and spatiotemporal neural dynamics of auditory information have also been characterized at millisecond resolution^48–50^.

When it comes to multisensory perception, high-order association areas such as superior temporal gyrus/sulcus (STG/STS) and intraparietal sulcus (IPS) are identified as multisensory integration regions^51,52^, because they respond to multimodal stimuli^53–55^ and even exhibit higher responses than unimodal input^56–58^. However, multisensory perception is not as simple as each type of sensory information being processed in its modality-specific brain regions and integrated in high-level association areas^59,60^. Numerous studies have shown that cross-modal interactions occur early in primary sensory regions with anatomical^61–65^, electrophysiological^64,66–69^ and neuroimaging^70,71^ evidence. Therefore, the entire brain can be regarded as multisensory^72–75^.

However, the functional roles of different brain areas in multisensory perception and the associated neural temporal dynamics remain less investigated, particularly in naturalistic contexts. Previous work on audiovisual perception often used illusions^76–80^ or rudimentary experimental stimuli like visual flash and auditory tones so that researchers could easily manipulate stimulus features such as temporal synchrony^81–84^, spatial location^85–88^, or modality reliability^89–92^ to examine multisensory integration^93,94^. Nevertheless, simple stimuli lack ecological relevance and cannot fully capture how humans perceive real-world audiovisual events. Some studies used image/sound pairs^95–99^ to investigate audiovisual perception, but most real-world audiovisual events do not remain visually still when dynamic sounds are produced. Recently, there has been growing evidence that naturalistic stimuli can better capture how real-world perception unfolds in the human brain^100^ and thus it is important to understand the neural basis underlying perception of naturalistic audiovisual events.

To address this, we employed naturalistic audio-video stimuli that capture common audiovisual events, and examined neural representations using multivariate pattern analysis on both functional magnetic resonance imaging (fMRI) and electroencephalogram (EEG) data. We used computational models to characterize how different types of stimuli information are processed across brain regions and over time. By teasing apart neural processes involved for distinct levels of information, we identified early asymmetrical cross-modal interaction: low-level acoustic information was represented in the early visual regions, but no low-level visual features were represented in the auditory areas. We also found that high-level categorical and semantic information emerged in multi-modal association areas, suggesting the late cross-modal interaction and its role in integrating conceptual information. We observed that visual and auditory features were processed with similar onset but distinct temporal dynamics, and high-level information was resolved later in time. Furthermore, we found a two-branch audio-video deep neural network model failed to capture early asymetrical cross-modal interaction, suggesting early fusion is an essential part to build a biologically plausible model of audiovisual neural processing. Lastly, we combined the fMRI and EEG responses through representational similarity framework^101^ to resolve neural activity in both space and time at the whole brain scale.

## 2 Results

To capture naturalistic audiovisual events, we curated 60 one-second videos with matching visuals and sounds. To exclude additional language or emotional processing, we chose only categories of animals, objects, and scenes with emotionally neutral contents (see Supplementary Table 1 for descriptions of each video). We recorded fMRI and EEG data separately while participants (N=22) passively viewed the videos with the accompanying sounds and performed an orthogonal oddball detection task to maintain their vigilance (Figure 1A). Each video was presented 11 times for fMRI and 12-15 times for EEG. The purpose of collecting both fMRI and EEG data is to combine the advantages of the two techniques and examine the brain responses with high spatial and temporal resolution.

**Figure 1:**
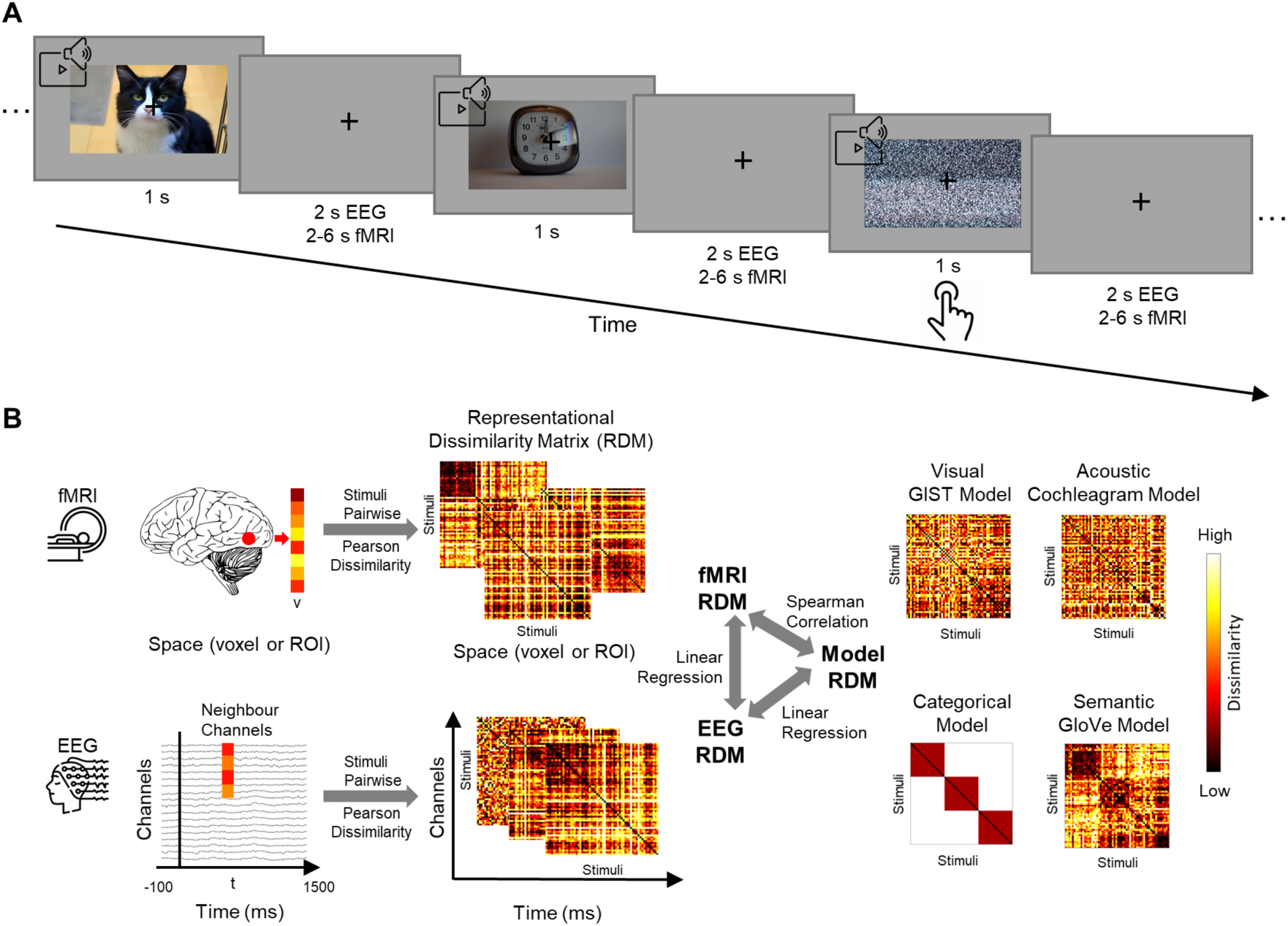
Experimental design and data analysis scheme. **(A)** The experiment consisted of two sessions in which fMRI and EEG data were recorded separately. In each run of the experiment, 60 one-second naturalistic audio-video stimuli were randomly presented with 10 noise videos in between. The participants (N=22) were instructed to view the videos with accompanying sounds. To maintain their vigilance, they were instructed to press the button when they detected the noise video. The data for the noise videos were excluded from the data analysis. **(B)** The stimuli-evoked fMRI estimates were extracted for a region of interest (ROI) or a voxel within a searchlight sphere (radius = 4 voxels). For each pair of stimuli, the dissimilarity between the fMRI patterns was calculated using Pearson correlation distance and stored in a symmetric representational dissimilarity matrix (RDM) indexed by the stimuli. Therefore, the size of the RDM is 60 x 60. The preprocessed EEG signals were extracted from −100 ms to 1,500 ms relative to stimulus onset. At each time point and for each channel, the signals with the neighboring channels were used to construct the RDM using the stimuli-pairwise Pearson correlation dissimilarity measure. As a result, neural representations were captured by fMRI RDMs in space and EEG RDMs in time. To assess how stimuli information was represented in the neural responses, different types of models were selected to characterize the visual (by GIST descriptors^102^), acoustic (by cochleagram model^103^), categorical (by predefined categories), and semantic (by GloVe word embeddings^104^) information. Each model RDM was created using the same stimuli pairwise Pearson distance measure on the model responses. Quantifying the correspondence between the model and neural RDMs by Spearman’s correlation or linear regression reflected how the model information was represented in the brain and processed over time. Calculating the correspondence between the fMRI and EEG RDMs resolved the neural activity at both high spatial and temporal resolution.

After data preprocessing, we applied multivariate pattern analysis and extracted the neural representations of the stimuli using representational similarity analysis^105^ (Figure 1B; see Methods). For fMRI, we obtained the evoked response patterns for each stimulus at either a region of interest (ROI) or a searchlight voxel sphere^106^ and constructed a representational dissimilarity matrix (RDM) by calculating the Pearson correlation distance between the fMRI patterns for each pair of stimuli. For EEG, instead of using a conventional approach to create a single RDM at each time point with whole-brain channels^35,36^, we created RDMs at each time point for each channel using its neighboring channels (i.e. a searchlight in EEG channel space). This method aims to preserve the EEG spatial information and prevent the visual information dominating the auditory information when all channels are treated equally.

To examine how different levels of stimuli information are represented in the brain, we used different computational models to capture low-level visual (GIST descriptors^102^) and acoustic features (Cochleagram model^103^) and high-level categorical and semantic features (GloVe word embeddings^104^) (Figure 1B). Model response patterns to each stimulus were extracted to create an RDM using the same Pearson distance metric. By calculating the similarity between neural and model representations, we aim to assess how the model information is represented in the brain or processed across time.

### 2.1 Asymmetrical neural representations of early cross-modal interaction in early visual regions

First, we aimed to investigate where visual and auditory information is represented in the brain during the perception of naturalistic audiovisual events using fMRI. To this end, we used GIST descriptors^102^ and the Cochleagram model^103^ to capture low-level visual and acoustic features respectively. We first examined information representation in atlas-defined visual- and auditory-related brain regions of interest (ROIs). We employed the Human Connectome Project multimodal parcellation (HCP-MMP) atlas^107^ and obtained neural representations in the form of ROI RDMs in visual regions including early, ventral, dorsal, and lateral visual areas, in auditory regions including early and association auditory areas, as well as in high-level multi-modal association areas (Figure 2A). We correlated the fMRI ROI RDMs with each model RDM using Spearman’s rank correlation. We partialed out the contributions of other tested models, because two-high level model RDMs showed slight but insignificant correlations with the visual feature model (categorical model: *ρ* = 0.039*, p* = 0.098; semantic model: *ρ* = 0.043*, p* = 0.071) and acoustic feature model (categorical model:*ρ* = 0.043*, p* = 0.067; semantic model: *ρ* = *−*0.034*, p* = 0.143). Notably, the visual and acoustic model RDMs did not correlate with each other (*ρ* = *−*0.030*, p* = 0.194). For statistical testing, we applied the sign permutation test (10,000 randomizations) with FDR correction at p<0.001.

**Figure 2:**
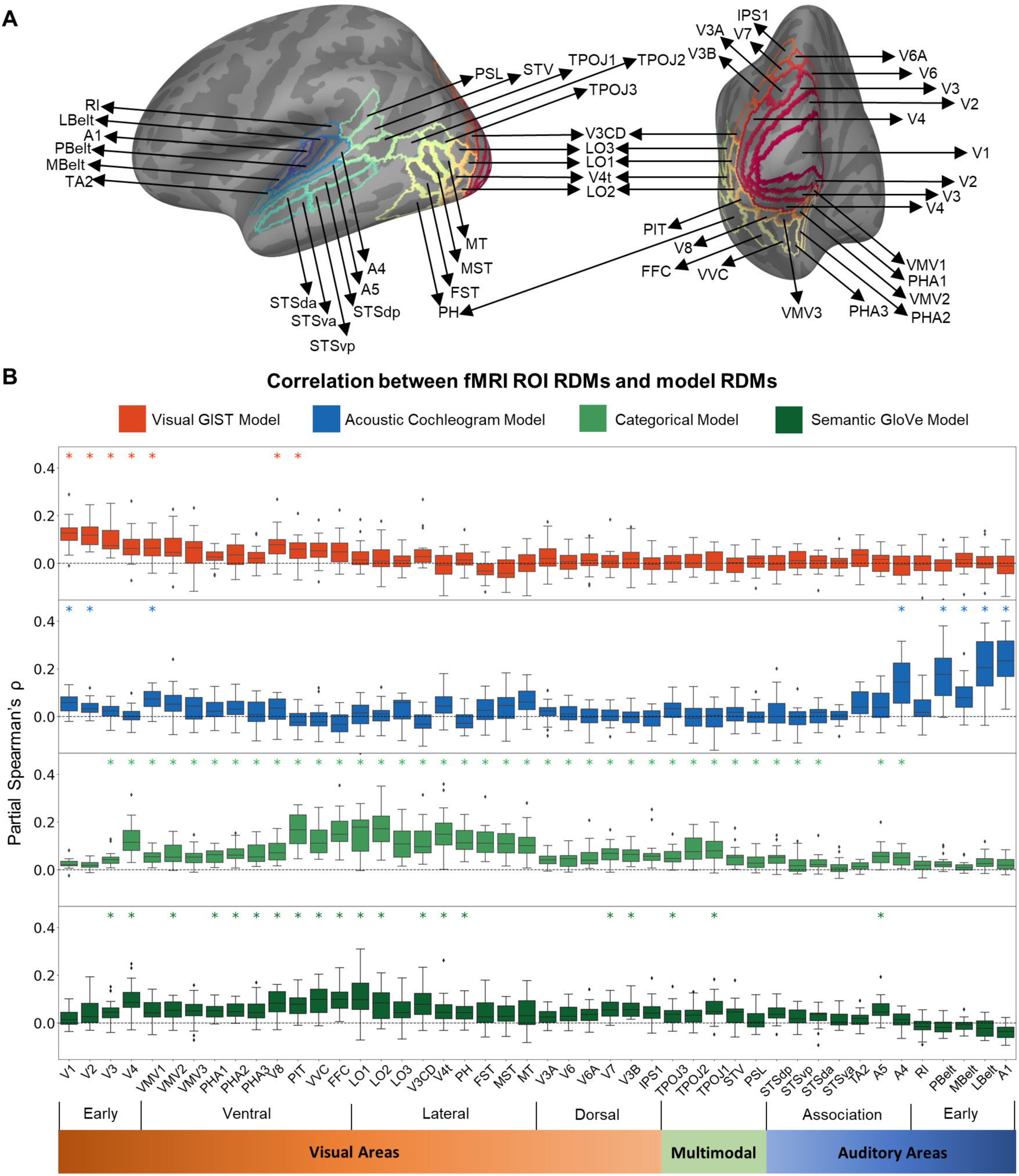
Comparisons between fMRI ROI RDMs and model RDMs. **(A)** Visualizations of the visual, auditory, and multi-modal brain regions selected from Human Connectome Project multimodal parcellation (HCP-MMP) atlas^107^. **(B)** The Spearman’s correlation between fMRI ROI RDMs and model RDMs with partialing out all other models (except for the correlation between categorical and semantic model because they are highly correlated with *ρ* = 0.480). The boxplots depict the correlation values across subjects for a given ROI denoted in the x-axis. Colors represent four different models. Asterisks above the boxplots indicate significant RDM correlations (sign permutation test with 10,000 randomizations, FDR-corrected at p<0.001).

The activity in the primary visual cortex (V1) best correlated with the low-level visual feature model, and the ROI-model RDM correlation decreased along the visual hierarchy from V1 to V4 (Figure 2B). Neural representations in some regions of the ventral visual stream showed smaller but significant correlations, including V8, the ventromedial visual 1 (VMV1), and the posterior inferotemporal complex (PIT). No significant correlation was observed in the dorsal and lateral visual regions or in the multimodal and auditory regions.

The low-level acoustic model correlated best with neural representations in the primary auditory cortex (A1), and the strength of the correlation decreased along the auditory hierarchy (Figure 2B). We did not observe significant correlations in auditory associations and multi-modal areas. However, we found that some visual regions, especially V1, showed a significant correlation with the low-level acoustic feature model. This suggests that auditory information is integrated early in V1. Surprisingly, this early cross-modal integration was only observed asymmetrically. Auditory regions do not represent any visual information.

To better assess the spatial distribution of effects, we applied searchlight analysis^106^ in which fMRI response patterns within a voxel sphere (radius=4 voxels) were used to compute searchlight RDMs, which were then correlated with the model RDMs using the same partial correlation approach, yielding three-dimensional correlation maps showing how the model information is represented in the whole brain (Figure 3). We performed 1,000 sign permutation test with cluster-based correction (cluster-definition threshold p<0.001, cluster threshold p<0.01). The results corroborated what we found with the ROI analysis. With the visual feature model, we mainly found significant correlations in early and ventral visual areas, and the higher correlation was mainly observed in early visual regions. No significant correlation was found in auditory regions. For the acoustic feature model, not only the auditory regions but also early visual and ventral regions showed significant correlation, demonstrating the asymmetrical neural representations of early cross-modal interaction.

**Figure 3:**
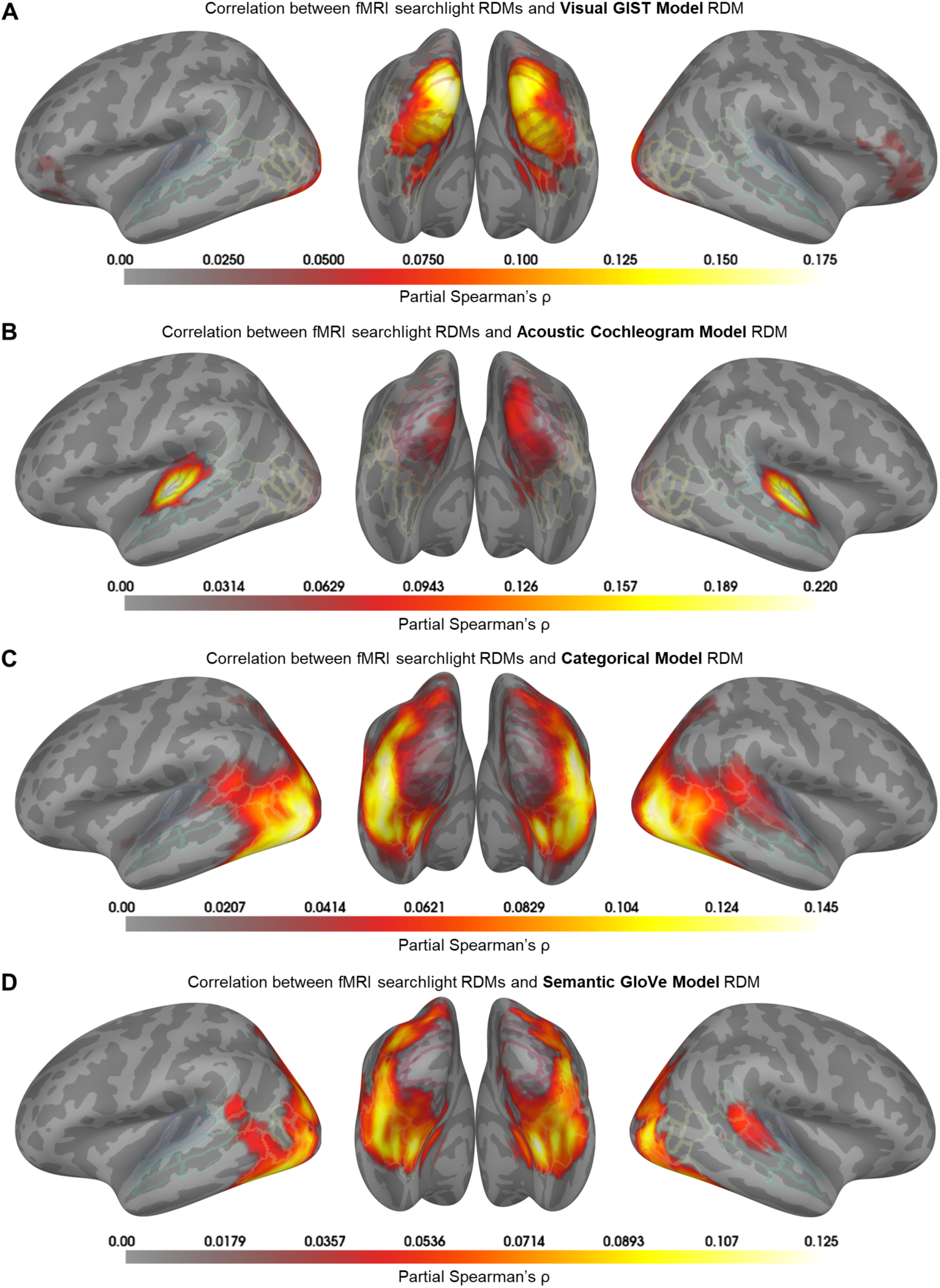
Comparisons between fMRI voxel searchlight RDMs and model RDMs. **(A)** Whole-brain correlation map with the visual GIST model. **(B)** Whole-brain correlation map with the acoustic cochleogram model. **(C)** Whole-brain correlation map with the categorical model. **(D)** Whole-brain correlation map with the semantic GloVe model. The correlation values are subject-averaged Spearman’s correlation between fMRI searchlight RDMs and model RDMs with partialing out all other models (except for the correlation between categorical and semantic model because they are highly correlated with *ρ* = 0.480). Only voxels with significant correlations are displayed and color-coded (cluster-corrected sign permutation test with 1,000 randomizations, cluster-definition threshold p<0.001, cluster threshold p<0.01).

### 2.2 Categorical and semantic representations in high-level visual, auditory and multi-modal regions

Next, we aimed to examine where in the brain high-level abstract information is represented. We first assessed categorical representations, using predefined categories to create categorical RDMs in which within-category dissimilarities were 0 and between-category dissimilarities were 1. To create a semantic RDM, we calculated pairwise Pearson distances between word embeddings from the GloVe model^104^ generated from text descriptions of the video contents. The categorical and semantic RDMs were highly correlated (*ρ* = 0.480*, p <* 0.001*^∗^*), so when comparing them with the fMRI RDMs we did not partial out the correlation between the two high-level models.

In the ROI analysis, we found that categorical information was represented in V3 and all high-level visual areas (Figure 2B). The high correlations were found in ventral regions such as the PIT, the ventral visual complex (VVC), and the fusiform face complex (FFC), as well as in lateral regions such as the three lateral occipital (LO) areas. Categorical information was not represented in early auditory areas but was present in high-order auditory regions such as A4, A5, and superior temporal sulcus (STS). All multi-modal association areas contained category information. These results suggest that high-level categorical information was resolved in the late stages of visual and auditory processing. A similar trend was observed for semantic representation, except that some regions showed lower or no significant correlations compared with the categorical model (Figure 2B).

The searchlight results (Figure 3) confirm that the categorical and semantic information was mainly represented in high-level visual, auditory, and multi-modal regions. This suggests that what is integrated in the established multi-modal association area is high-level conceptual information of multi-modal stimuli.

### 2.3 A two-branch audiovisual deep neural network failed to capture the early cross-modal interaction

Next, we asked whether the observed early asymmetric cross-modal interaction in the brain could emerge as a result of high-level cross-modal representation matching of visual and auditory cortices. To this end, we chose a pre-trained deep neural network model^108^ composed of two separate streams which receive either video frames or audio spectrograms, similar to the brain’s separate visual and auditory pathways (Figure 4A). However, the two model branches lack direct connections, unlike the brain with projections between visual and auditory areas. The model was trained on AudioSet, a set of naturalistic audiovisual event stimuli^109^, with a contrastive learning framework^110,111^ to match the video and audio representations for the same stimulus and repel the representations for different stimuli. Compared with supervised training, contrastive training allowed the model to learn better representations and achieve higher task performance^108^. To evaluate the hierarchical correspondence between the model and the brain, we extracted model activations from seven blocks of each model stream and used them to construct RDMs.

**Figure 4:**
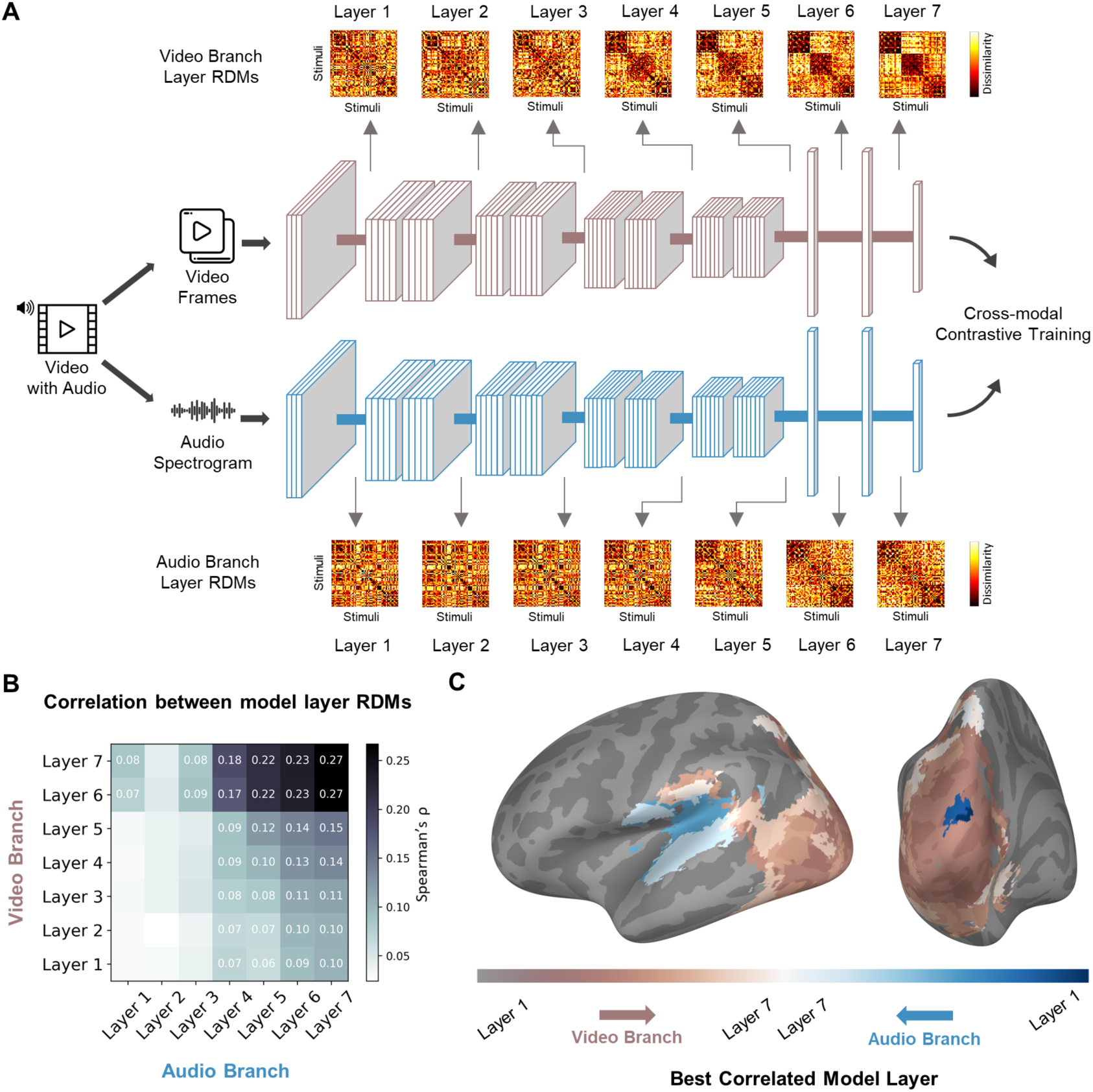
Comparisons between fMRI voxel searchlight RDMs and two-branch deep neural network (DNN) pretrained on audio-video stimuli. **(A)** Schematic illustration of the DNN model^108^ trained on audio-video stimuli^109^ with two separate branches receiving video frames and audio spectrogram respectively. The outputs of two streams were trained to match for the same video and repelled for different videos through a contrastive learning framework^110,111^. The model activations of experimental video stimuli were extracted at different blocks of each model branch to create an RDM using the stimuli pairwise Pearson distance measure, yielding a total of 14 RDMs to capture the representations of 7 hierarchical layers of each model stream. **(B)** The Spearman’s correlation values between each model layer RDMs. Only significant correlations were displayed numerically (10,000 paired permutation testing, FDR-corrected at p<0.01). **(C)** The best-correlated layers were mapped on the brain after correlating each DNN layer RDM with each voxel RDM. The color indicates the layer of the model that showed the highest correlation (cluster-corrected sign permutation test with 1,000 randomizations, cluster-definition threshold p<0.001, cluster threshold p<0.01).

We observed that representations in early layers of video and audio branches did not exhibit any significant correlations, but representations in higher layers did show significant correlations and the values increased with layers (10,000 paired permutation testing, FDR-corrected at p<0.01; Figure 4B). This confirms that the model gradually learned to match high-level representations. Comparing the layer RDMs of the DNN model with the feature model RDMs showed that the model did not exhibit early-cross model interaction as observed with human fMRI responses (see Supplementary Figure 1). We next applied searchlight analysis to identify the brain voxels that showed a significant correlation with model layer RDMs (cluster-corrected 1,000 sign permutation test, cluster-definition threshold p<0.001, cluster threshold p<0.01; Figure 4C). The DNN model showed significant correlations with the visual, auditory, and multimodal regions. We observed that the model exhibited modality correspondence and hierarchical progression: early layers correlated with early brain regions, while later layers correlated with higher-level regions. However, some voxels in early visual areas correlated best with early layers of the audio branch, again demonstrating the early asymmetric cross-modal interaction in the brain is absent between the early layers of the model branches. This implies that including an early integration component is needed to build a biologically plausible computational model of audiovisual neural processing.

### 2.4 Temporal dynamics of audiovisual information representations

Next, we aimed to characterize how neural representations of different stimuli information unfold over time using EEG data. We did not adopt the conventional method of creating time-resolved EEG RDMs by treating all channels equally^35,36^ because the larger size of the visual cortex and its higher signal-to-noise ratio may overshadow the information captured from the auditory cortex in EEG recordings. Instead, we computed time-resolved EEG RDMs using a searchlight approach (Figure 1B; see Methods). We used the same stimuli pairwise Pearson dissimilarity metric and constructed 64 channel-wise searchlight RDMs for each millisecond from -100 to 1,500 ms relative to stimulus onset. At each time point, we then fitted a non-negative linear regression to evaluate how much variance in the model RDM can be explained by the 64 EEG channel RDMs. This approach allowed the linear regression model to learn the positive weights for each channel instead of assuming equal weights. As a result, we acquired a time course of explained variance for each model and subject (adjusted *R*^2^; see Methods), which quantified how much the model information was represented in the neural responses at each time point. To estimate the onset latencies, we applied jackknife-based approach^112–114(see^ Methods). To estimate the peak latencies, we performed 10,000 bootstrapping to estimate the confidence interval.

Figure 5 shows the time course of the adjusted explained variance *R*^2^ for four different models and the significant time windows. For the low-level visual feature model, the *R*^2^ became significant at 64 ms (95% confidence interval 60-69 ms), and first peaked at 96 ms (85-104 ms). The low-level acoustic feature representation emerged at 67 ms (61-72 ms) and first peaked at 86 ms (74-94 ms). Both the onset and first peak latencies did not differ statistically (onset: *p* = 0.509; first peak: *p* = 0.690), suggesting that visual and acoustic information were processed almost simultaneously. However, the maximum peak (i.e. global peak) for the auditory model was at 224 ms (180-295 ms), later than the maximum peak of the visual model at 134 ms (97-199 ms) (*p <* 0.001), implying that the extraction of salient information from sounds may require more accumulated time than visual scenes. Categorical information emerged later, at 169 ms (142-195 ms), with a peak at 205 ms (183-230 ms). Semantic representation was resolved with similar timing, with onset at 163 (134-192) ms and peak at 227 (181-258) ms. No significant difference was found between the two high-level models for either onset (*p* = 0.158) or peak latencies (*p* = 0.774).

**Figure 5:**
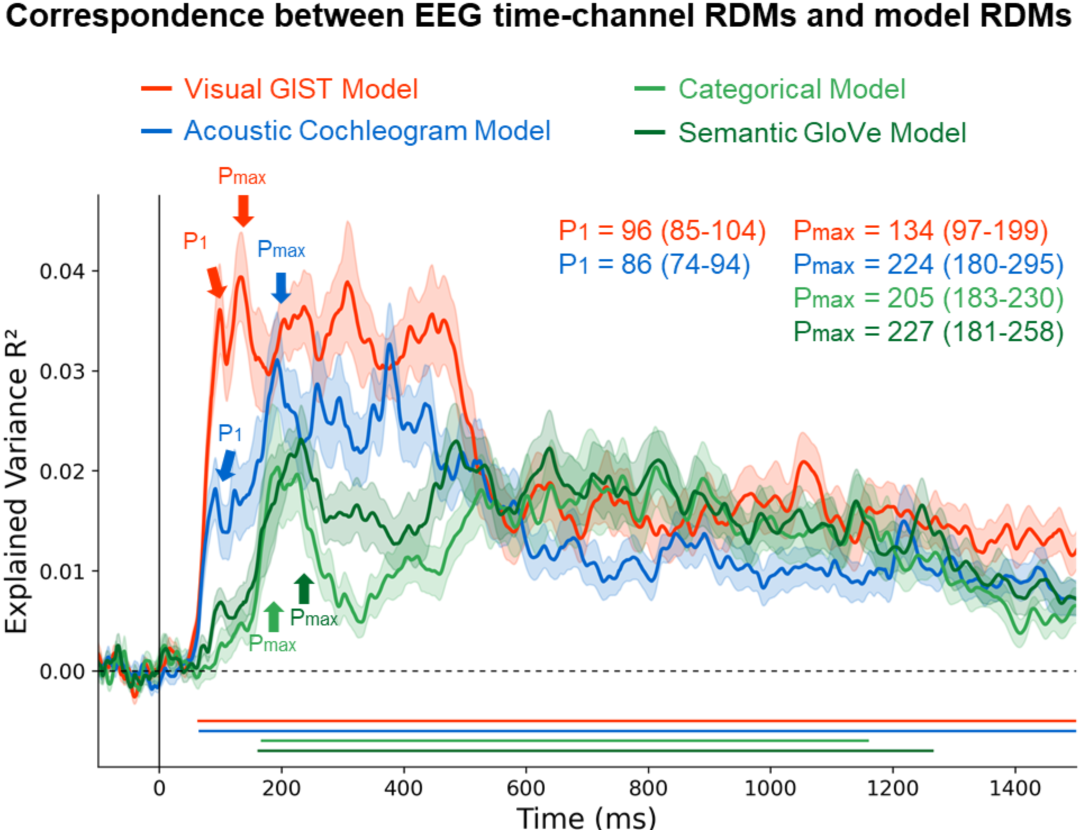
Correspondence between EEG time-channel RDMs and model RDMs. Time course of adjusted explained variance when EEG channel RDMs were fitted using non-negative linear regression to predict the model RDMs (see Methods). Each color-coded curve represents the subject-averaged adjusted explained variance *R*^2^ values for a given tested model. The shaded areas denote the stand error of the mean. The bottom lines indicate the significant time points at which jackknifed subsamples passed the jackknife-based baseline criteria (see Methods). The color-coded arrows denote peak latencies for a given tested model, with corresponding values and 95% confidence intervals determined by 10,000 bootstrapping subject samples.

### 2.5 Spatiotemporally resolved audiovisual processing

Finally, we aimed to resolve neural activity with high spatial and temporal resolution to characterize audiovisual processing in both space and time. To this end, we fused the fMRI and EEG RDMs, inspired by the similarity-based fMRI-M/EEG fusion^38,41,101^. To benefit from the spatial resolution available from EEG channels, we extended the fMRI EEG fusion technique by fitting non-negative linear regression on the EEG channel RDMs to predict each voxel RDM, with the adjusted *R*^2^ as the measure of EEG-fMRI correspondence. As in Figure 6, significant correspondence first emerged in early auditory and visual regions 60–65 ms after stimulus presentation, and then spread to high-level regions, consistent with hierarchical processing in both vision and audition^4–6^.

**Figure 6:**
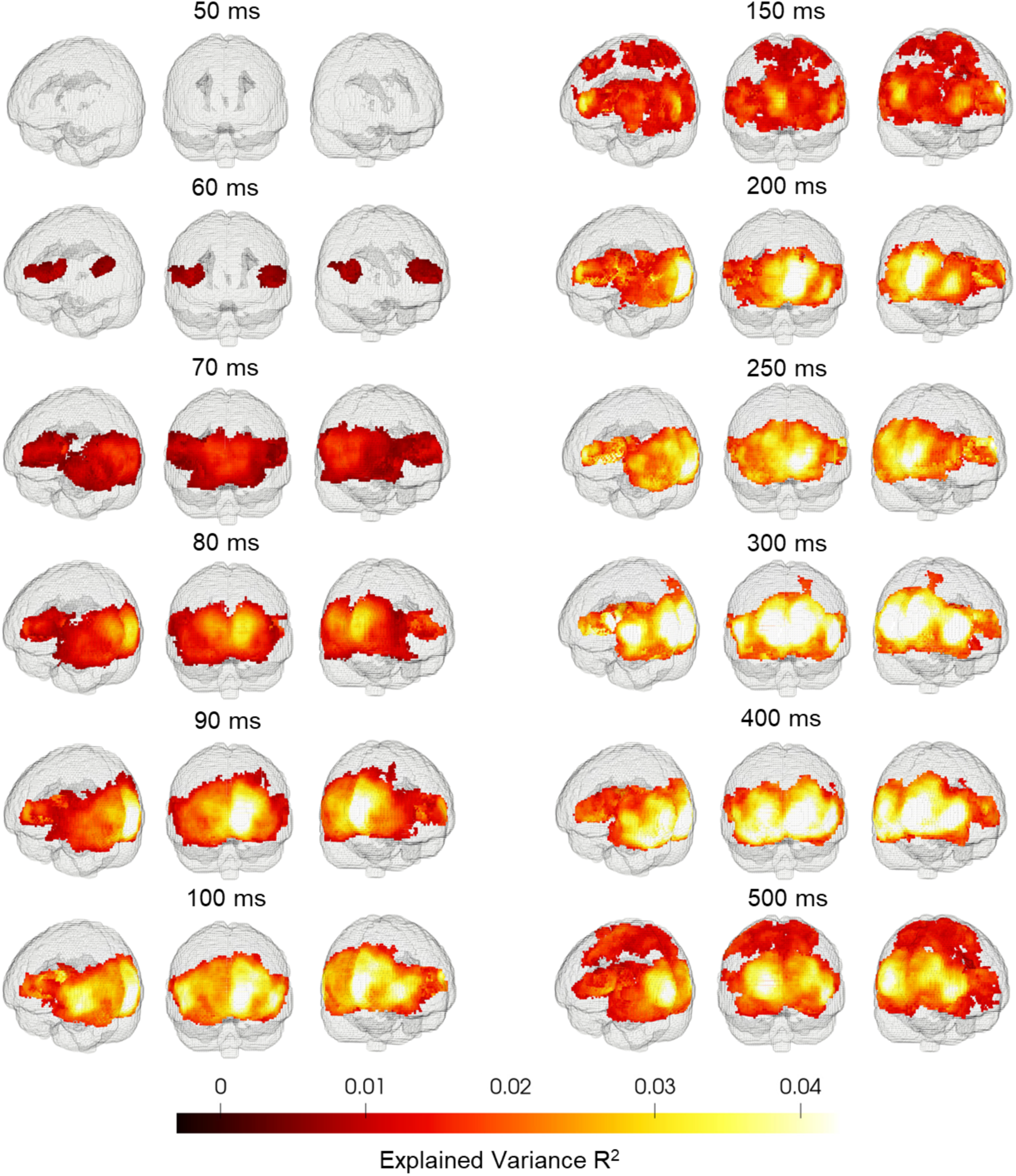
Fusion of fMRI voxel searchlight RDMs and EEG time-channel RDMs. Wholebrain maps of subject-averaged adjusted explained variance scores when EEG channel RDMs were fitted using non-negative linear regression to predict the fMRI searchlight RDMs for selected time points (cluster-corrected sign permutation test with 1,000 randomizations, cluster-definition threshold p<0.001, cluster threshold p<0.01). For results of all time points, see the Supplementary Movie.

We also examined ROI-specific temporal profiles. Figure 7A shows the correspondence time course for selected visual and auditory regions. We found that the onset latency for V1 was at 64 ms (60-69 ms) and for A1 was at 62 ms (59-65 ms), with no significant difference (*p* = 0.380). This suggested that V1 and AI were activated with similar processing speed. The first peak latencies for V1 and A1 showed no significant difference (*p* = 0.949), with activity in V1 first peaked at 100 ms (94-105 ms) and in A1 first peaked at 91 ms (81-110 ms). However, the maximum peak latencies differed significantly (*p <* 0.001) with V1 at 124 ms (96-130 ms) and A1 at 224 ms (191-372 ms). When quantifying the maximum peak time for each ROI with bootstrapping, we found evidence for a temporal hierarchy (Figure 7B), in which high-level brain regions take more time to reach the peak than low-level regions. Taken together, by computing the correspondence between fMRI and EEG RDMs, we provided a spatiotemporally resolved view of neural activity during the perception of naturalistic audiovisual events.

**Figure 7:**
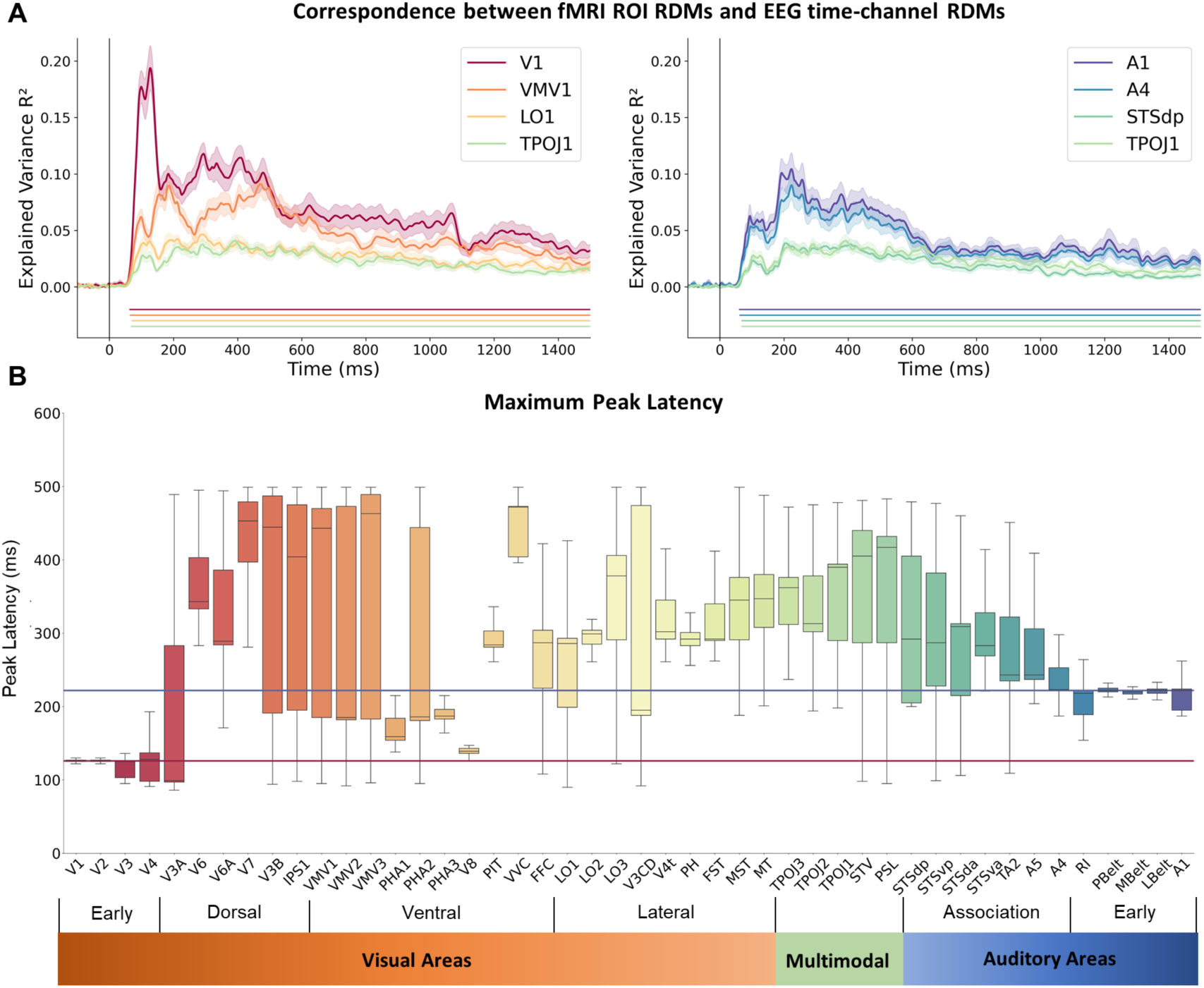
Correspondence between fMRI ROI RDMs and EEG time-channel RDMs. **(A)** Time course of adjusted explained variance when EEG channel RDMs were fitted using non-negative linear regression to predict the fMRI ROI RDMs for selected brain regions. Each color-coded curve represents the subject-averaged adjusted explained variance *R*^2^ values with the shaded areas denoting the stand error of the mean for the given ROI. The bottom lines indicate the significant time points at which jackknifed subsamples passed the jackknife-based baseline criteria (see Methods). **(B)** The boxplot of the maximum peak latency of the adjusted explained variance over 10,000 bootstrapping resamples when EEG channel RDMs were fitted using non-negative linear regression to predict the fMRI ROI RDMs.

## 3 Discussion

Using naturalistic audio-video stimuli, we investigated neural responses during the perception of real-world audio-visual events recorded by fMRI and EEG. We examined which brain regions represented low-level visual and acoustic features and high-level categorical and semantic information as well as when they were processed. By examining neural processes involved for different types of information, our results revealed both early and late cross-modal interactions, with their associated brain areas, temporal dynamics, and functional roles. We further found that early cross-modal interaction did not emerge in a two-branch deep neural network model, suggesting that early fusion is needed to build a biologically plausible model of audiovisual perception. Finally, through the fusion of representational information captured by fMRI and EEG, we resolved neural activity with high spatial and temporal resolution.

Previous findings show that cross-modal interactions occur at different processing stages and can be observed as early as primary sensory regions^73–75^. However, what roles these interactions play and how their functions differ in each brain region remains unclear, especially under more naturalistic settings. Here we found that during the perception of naturalistic audiovisual events, the early visual areas represented low-level acoustic information in addition to low-level visual features, suggesting early cross-modal interaction. This can be supported by the fact that primary visual cortex receives direct projection from the auditory cortex^115,116^, with evidence from animal trace and electrophysiology studies^117,118^ as well as human functional and structural connectivity^119–121^. Previous work^122,123^ showed that when participants heard natural sounds only, the content of sounds can be decoded in the V1 in both healthy and blind populations, but whether such auditory information comes for feedback projection from the primary auditory cortex or high-level extrastriate and multisensory areas remains unknown. Here we demonstrated that the auditory information V1 contains is mainly low-level acoustic features instead of high-level concepts, suggesting the feedback signals mainly come from the early auditory cortex, for its role in processing low-level acoustic features^124–126^.

However, we did not find any visual information in the auditory cortex. Previous studies found that the activation of the auditory cortex is modulated by additional visual input^71,96,127–129^. Nevertheless, it does not mean that our results are inconsistent because our study can not examine the modulation effect, which can only be observed when we compare the perception of audiovisual stimuli with audio stimuli alone. Our results show that at the multivariate representational level, the cross-modal interaction in low-level feature representations is not bidirectional. Instead of combining low-level visual and acoustic information in both sites, the brain chose the visual cortex to integrate the acoustic features and not otherwise. One benefit of such unidirectional integration could be to reduce the redundant load of bidirectional communications of multi-modal features. However, the underlying mechanisms still need further exploration.

Our further analyses showed that established multi-modal association regions represent high-level categorical and semantic information. Together, our findings suggest that there exist at least two different stages of cross-modal interactions with distinct roles: (1) Early crossmodal interaction in early visual regions. The acoustic features processed through the early auditory areas are projected to early visual areas. One possible purpose of this process could be to ensure a consistent multi-modal percept of the objects^75^. If two modality systems work in complete isolation and only merge final high-level outputs, it is likely that the final outputs may be mismatched (e.g., when seeing a house and hearing a dog). Therefore, early fusion may help solve potential mismatching problems. (2) Late cross-modal interaction in multi-modal regions. The high-level conceptual information extracted from each modality converges in these regions. The integration process here may require further computations and thus result in the super-additive activations observed in previous studies^56,57,130,131^. Future work with semantically mismatching multi-modal stimuli can violate the high-level convergence and may potentially help to investigate the mechanisms of different integration processes during multisensory perception.

Currently, deep neural network (DNN) models serve as the best models of human visual or auditory system^132–136^. However, their similarity with human brain responses in multisensory perception is less explored^137^. Generally, the match between DNN models and the brain depends on multiple factors, such as the training dataset, the objective function, and the model architecture^136,138,139^. Here we evaluated a pretrained DNN model^108^ with a two-separate-branch architecture trained on a naturalistic audio-video event dataset to learn the high-level audio-visual representational correspondence^140,141^. We found alignment in hierarchical progression between the model and the brain, but the model did not capture early cross-modal interaction. One possible reason is that the model only has high-level integration at the end of each stream and lacks early connections between the two branches, unlike the brain which has projections between the early visual and auditory cortices^61,63–65^. Therefore, including the early-fusion component is needed to build a biologically plausible model. Some recent computational work has implemented the idea of early cross-modal interaction, either unifying the audio and visual branches^142^ or adding a branch of early fusion^143^, and achieved enhanced performance in related audiovisual tasks. Future work can examine whether models with early fusion might show better correspondence with the human brain, and whether they can elucidate the computational mechanisms of different cross-modal interactions.

Previous studies have revealed the temporal neural dynamics during the perception of natural images^35,38^ or sounds^34,49,50^. However, the temporal dynamics during multisensory perception of naturalistic stimuli remain less investigated. To our knowledge, no study to date has investigated whether any disparities exist in the processing speed of different sensory modalities during multisensory perception. Here we found that the neural representations of visual and acoustic features emerged and first peaked almost simultaneously with no significant differences. A similar trend was also observed for neural activity in V1 and A1, suggesting that the visual and auditory information reach and activate the primary sensory regions with similar speed. However, the maximum peak time differed for the two sensory modalities, with the maximum peak time delayed for A1 (or acoustic features) compared with V1 (or visual features). One possible explanation could come from the inherent distinctions between the two modalities. Unlike visual scenes, sounds involve temporal accumulations, particularly for natural stimuli. We observed that high-level categorical and semantic information was resolved at late stages, consistent with previous findings showing that high-level conceptual information is processed later than low-level stimulus features^35,36,49^. However, our results cannot tease apart when acoustic information is processed in A1 or V1 given that early cross-modal interaction involves acoustic features represented in both areas. Future work is needed to address this with different experimental designs or analysis approaches. As for the late cross-modal interaction, we can infer the timing by the active time of multimodal association regions or high-level information representations.

Multisensory integration requires the brain to solve the question of whether the incoming sensory information of distinct modalities comes from the same source. A proposed mechanism is Bayesian causal inference^144–146^ and many studies have examined the neural basis of this process^90,92,147^. They mainly relied on simple, experimentally controlled stimuli and focused on the processing components during causal inference. In our study, we assumed that integration occurs naturally with naturalistic audiovisual stimuli, in which we did not manipulate the modality reliability or the temporal coherence. When curating the stimuli, we ensured minimal effort was required to infer whether the visuals or sounds were matched. We mainly focused on how different types of stimulus information are resolved in the brain and across time, thus characterizing multisensory perception with more ecological validity. Nevertheless, we encourage future studies to examine the neural spatiotemporal dynamics of the Bayesian causal inference process under more naturalistic settings.

## 4 Methods

### Participants

Twenty-two subjects participated in this study (12 females; mean age=23.04, SD=3.25). All participants were right-handed with normal or corrected-to-normal vision and normal hearing. They reported no history of neurological or psychiatric disorders and provided written consent. The study was approved by the Non-Medical Research Ethics Board at Western University and all participants were compensated for their time.

### Stimuli

We curated 20 naturalistic videos for each category of daily common objects, animals, and scenes. Each video includes representative visuals and sounds of a certain event. These naturalistic video stimuli were found online and cropped into one second with a frame rate of 25 per second and a frame size of 1920 x 1080. We made sure the videos have semantically matched high quality audios. The audios were root-mean-squared normalized and resampled to 48 kHz.

### Procedure

The experiment consisted of two sessions with the same task but recorded with fMRI and EEG separately. The fMRI session includes 11 runs and the EEG session includes 12-15 runs. In each run, all video stimuli were presented in random order with 10 noise videos in between. Each video was presented at the center of the screen (view angle *≈* 5°) with a black fixation cross. The fMRI and EEG sessions differed in the inter stimuli interval (ISI), with a 2 second fixed ISI for EEG and a 2-6 second jittered ISI for fMRI. The task of the subjects was to view the videos and fixate at the center of the screen. Once they detected the noise video, they were instructed to press a button, and the purpose of this task was to ensure their vigilance. The participants were exposed to all stimuli before the data collection and no category information was provided.

### EEG data acquisition and preprocessing

EEG data were recorded using Biosemi 64-channel system with a sampling rate of 2048 Hz. An optic sensor was attached to the screen to ensure the onset of stimuli presentation matched with EEG stimuli event markers. The data were preprocessed using FieldTrip^148^. Bad channels were removed and re-reference was applied using the common average reference. The data trials were epoched from -100 to 1,500 ms relative to the stimuli onset, demeaned using the average value of the 100 ms prestimulus period, and high-pass filtered at 0.1 Hz. Trials with jump and muscle artifacts were detected and removed using the FiedlTrip automatic artifact rejection method and trials with high variance were visually detected and removed. Eye blink and movement artifacts were identified and removed using independent component analysis (ICA). Finally, the data were low-pass filtered at 30 Hz, resampled at 1000 Hz, and normalized with multivariate noise normalization^149^.

### fMRI data acquisition

Functional MRI data were acquired using a Siemens 3T Magnetom Prisma Fit MRI scanner at the Centre for Functional and Metabolic Mapping at Western University. T2*-weighted, singleshot, gradient-echo echo-planar imaging (GE-EPI) sequence was used with the following parameters: TR = 1000 ms, TE = 30 ms, flip angle = 48°, FOV = 210 mm, multiband factor =5, slice thickness = 2.5 mm, voxel size = 2.5 × 2.5 × 2.5 mm, and 60 axial slices covering the whole brain. High-resolution anatomical images were also acquired using a T1-weighted MPRAGE sequence with the following parameters: TR = 2300 ms, TE = 2.98 ms, flip angle = 9°, FOV = 256 mm, slice thickness = 1.0 mm, voxel size = 1.0 × 1.0 × 1.0 mm, and 192 sagittal slices.

### fMRI data preprocessing and analysis

fMRI data were preprocessed using the fmriprep pipeline^150^ (version 20.2.5), an open-source tool developed for automated preprocessing of fMRI data. A series of standard preprocessing steps were applied with the default settings and achieved through a combination of ANTs (http://stnava.github.io/ANTs/), FSL^151^, and FreeSurfer (http://surfer.nmr.mgh.harvard.edu/).

Specifically, T1-weighted structural images underwent brain extraction, tissue segmentation, and spatial normalization to the MNI152NLin2009cAsym template. Functional images were slice-timing corrected, head motion corrected, susceptibility-derived distortion corrected, coregistered to T1-weighted images, and resampled to the standard space. No spatial smoothing was applied.

Generalized linear modeling (GLM) was implemented using SPM 12 (https://www.fil.ion.ucl.ac.uk/spm/software/spm12/). We concatenated fMRI data across all runs and modeled the stimuli-evoked responses with the onsets and durations of each stimulus convolved with the canonical hemodynamic response function as the regressors. Movement parameters and run regressors were included as regressors of no interest. The SPM default 128-second high-pass filter was applied before GLM. For each of the 60 stimuli, we obtained a single parameter estimate and converted it into a t-value by contrasting it against the implicitly modeled baseline.

### fMRI and EEG multivariate pattern analysis

We followed representational similarity analysis^105^ to convert fMRI and EEG responses to representational space by constructing representational dissimilarity matrices (RDMs). The RDM is indexed by stimuli and each element value quantifies the dissimilarity between the measured neural responses to a pair of stimuli.

For fMRI regions of interest (ROI) method, we selected brain regions that are defined as visual, auditory, or multi-modal areas from Human Connectome Project multimodal parcellation (HCP-MMP) atlas^107^. For each ROI, we extracted fMRI activation patterns and created an RDM by calculating the 1-Pearson correlation between the fMRI patterns for each pair of stimuli. The value in the RDM is in the range of 0 to 2, where 0 denotes complete similarity and 2 denotes complete dissimilarity. For fMRI searchlight approach, we extracted the fMRI activity responses at each voxel using its local sphere voxels with a radius of 4 voxels (10 millimeters) and constructed an RDM again using the stimuli-wise Pearson correlation distance. This approach yielded 121,799 RDMs.

For EEG data, we did not adopt the conventional method to use the activity patterns of whole-brain channels to create a single RDM at each time point. The shortcoming of this approach is that it treats all channels equally to convert the EEG responses to representational space. When visuals and sounds are processed simultaneously, because the visual cortex is much larger than the auditory cortex, thus more channels are devoted to visual areas and then the RDM created is biased toward the visual information. To address this, we followed the idea of searchlight analysis and created one RDM for each EEG channel using its neighboring channels at each time point. As a result, we can preserve both the spatial and temporal information in the EEG signals, without obscuring the channel-wise information by simple average. Here we applied the same Pearson distance measure to calculate the dissimilarity values between the trial-averaged EEG signals to each pair of stimuli. This new approach yielded 64 RDMs at each millisecond from -100 to 1,500 ms, a total of 134,464 RDMs.

### Computational Models

We used GIST descriptors to capture the low-level visual features in the videos because the GIST descriptors summarize spectral and coarsely localized visual information, which are sufficient to identify the scene in an image^102^. To apply GIST descriptors, each frame image was convolved with 32 Gabor filters at 4 scales and 8 orientations to generate 32 feature maps of the same size as the input image. Then each feature map was divided into 16 regions with 4 x 4 grids and the feature values within each region were averaged, yielding a total of 521 (16 x 32) descriptors. The distance of the output values of descriptors between each pair of stimuli was calculated with Pearson correlation distance measure and stored in an RDM, which quantifies stimuli-wise representations of low-level visual features. After constructing 25 RDMs for each frame, we averaged RDMs across frames to obtain a single RDM.

To capture the low-level acoustic features, we used cochleagram model^103^ following^125^. The audio waveform of each video sound was convolved with 120 bandpass filters, which were spaced equally over an equivalent rectangular bandwidth (ERB) scale between 20 Hz and 10 kHz. The filter outputs were Hilbert transformed, raised to the 0.3 power, and downsampled to 400 Hz after applying an anti-aliasing filter. The frequency axis was transformed from an ERB-scale to a logarithmic frequency scale with 217 channels. The final output size for each sound was 217 x 400 and then the cochleagram features were vectorized and used to construct an RDM by computing the stimuli pairwise Pearson dissimilarity values.

To represent the high-level information of video stimuli, we used both category information and semantic embeddings extracted from the video text descriptions. The categories were predefined and guided to search for videos. The categorical RDM was created in which the dissimilarity value for a pair of stimuli from the same category was assigned 0 and from different categories was assigned 1. To extract semantic representations, we used pretrained GloVe (Global Vectors for Word Representation) model^104^, which is trained over a large corpus of text in an unsupervised manner and can effectively measure the semantic distance between a word pair. For each description of a video, the embedding vectors were extracted for each word and averaged to construct an RDM by measuring the Person correlation distance between each pair of stimuli.

To assess the similarity between the deep neural network model and the brain, we selected a pretrained two-stream model trained on audiovideo dataset AudioSet^109^ with a contrastive learning framework^110,111^ to match the video and audio representations for the same stimulus and repel the representations for different stimuli. The model architecture is similar to the brain structure, in which two model streams are fed with video or audio respectively, similar to separate visual and auditory pathways of the brain. To evaluate hierarchical correspondence, we extracted the model activations at the end of each seven blocks of the model streams and constructed the RDM with stimuli pairwise Pearson distance measure, yielding a total of 14 RDMs.

### Representational Similarity Analysis and EEG-fMRI Fusion Analysis

To quantify how representational information captured in the models is represented in the brain areas, we correlated model RDMs with fMRI ROI or searchlight RDMs with partial Spearman’s rank correlation. To assess how the model information unfolds over time, we fitted a nonnegative linear regression using 64 channel RDMs to predict the model RDM at each millisecond from -100 to 1,500 ms with respect to the stimuli onset, with the coefficient parameters enforced to be positive. Then we used the explained variance *R*^2^ as the measure to quantify the correspondence between the EEG channel RDMs and model RDMs. Because *R*^2^ increases with the number of predictors, we adjusted by averaging the *R*^2^ values when the channel RDMs were used to fit the random noise for 1000 randomizations and used this value as the baseline to correct the *R*^2^ scores when fitted to the model RDMs. We did not employ the adjusted *R*^2^ formula because it only applies to vanilla linear regression but not non-negative linear regression. To resolve the spatiotemporal dynamics of neural representation, we followed the idea of similarity-based M/EEG-fMRI fusion analysis^38,41,101^ but here at each time point, we fitted a non-negative linear regression using 64 channel RDMs to predict the fMRI ROI or searchlight RDMs and similarly used the adjusted *R*^2^ as the measure of correspondence between EEG and fMRI.

### Statistical Analysis

When we performed statistical testing on correlation values between fMRI RDMs and model RDMs, we applied non-parametric sign permutation, in which the signs of the data were randomly shuffled to create a null distribution and to compute the statistics against zero. For the results of the ROI analysis, we used FDR correction for multiple comparisons and applied threshold p < 0.001. For searchlight results, we applied cluster-level correction^152^, which takes into account the spatial proximity. The clusters were first determined by spatial contiguity at the cluster-definition threshold, and then a distribution of maximum cluster size was computed over permutations, and finally the clusters passing the cluster threshold were reported as significant. We chose the cluster-definition threshold p < 0.001 and cluster threshold p < 0.01.

To test whether correlations between each model layer RDMs were significant, we applied the paired permutation testing. For each pair of RDMs, we shuffled the order of values in one RDM and correlated with the left RDM using Spearman’s rank correlation. We repeated this process 10,000 times to create the null distribution and obtained p-values based on the null distribution. To address multiple comparisons, we applied FDR correction with a threshold of 0.01.

For EEG results, we applied jackknife-based method^112,113^ to estimate significant time points. We first calculated the standard deviation of the baseline values between −100 and 0 ms across all 22 subjects to establish the baseline criteria. Next, we compute 22 subsamples by excluding one subject each time and averaging the remaining subjects. For each of the jackknifed subsamples, a time point was considered significant only when the value at this time point was two times larger than the baseline standard deviation and the average value of the next 10 windows of 50 ms was also two times larger than the baseline standard deviation. Because the jackknifed subsamples were not independent, when computing subsequent statistics, we adjusted the variance by multiplying by a factor of (n-1) where n is the number of subjects. We computed the 95% confidence interval of onset latencies by multiplying the adjusted standard error by the corresponding t-statistic. To test differences in onset latencies for different models, we ran a two-sided paired *t* test. To estimate the 95% confidence intervals for peak latencies, we created 10,000 bootstrapped samples by sampling the subjects with replacement. To test differences in peak latencies for different models, we computed p-values using the paired differences of peak latencies from 10,000 bootstrapped samples.

For searchlight EEG-fMRI fusion results, we again performed non-parametric sign permutation test. However, because *R*^2^ values are all above zero, we tested against the value of the last 5% in the distribution instead of zero. we again applied cluster-level correction with cluster-definition threshold p < 0.001 and cluster threshold p < 0.01. For ROI-specific EEG-fMRI fusion results, we applied the same jackknifed-based approach to determine the significant time points and used bootstrapping method to compute the 95% confidence intervals of peak latencies.

## Acknowledgements

This study was supported by the Canada First Research Excellence Fund (CFREF) through a BrainsCAN grant to Y.M., a Vector Institute Research Grant to Y.M, and an NSERC Discovery Grant to Y.M.. The authors would like to thank Diana Dima for helpful comments and discussions on the manuscript.

## Author Contributions

Conceptualization: Y.H. and Y.M.; Methodology: Y.H. and Y.M.; Software: Y.H.; Validation: Y.H.; Formal Analysis: Y.H.; Investigation: Y.H.; Resources: Y.M.; Data Curation: Y.H.; Writing – Original Draft: Y.H.; Writing – Review and Editing: Y.H and Y.M.; Visualization: Y.H.; Supervision: Y.M.; Project Administration: Y.M.; Funding Acquisition: Y.M.

## Supplementary Information

### Supplemental Movie

The whole-brain EEG-fMRI fusion movies can be viewed via https://www.youtube.com/watch? v=8nc9BnuTosQ. It provided a spatiotemporally resolved view of whole-brain activity at each millisecond. The color-coded values denote correspondences between fMRI and EEG responses in representational space^101^ (1,000 cluster-corrected sign permutation test, cluster-definition threshold p<0.001, cluster threshold p<0.01).

## Supplemental Figures

**Figure S1:**
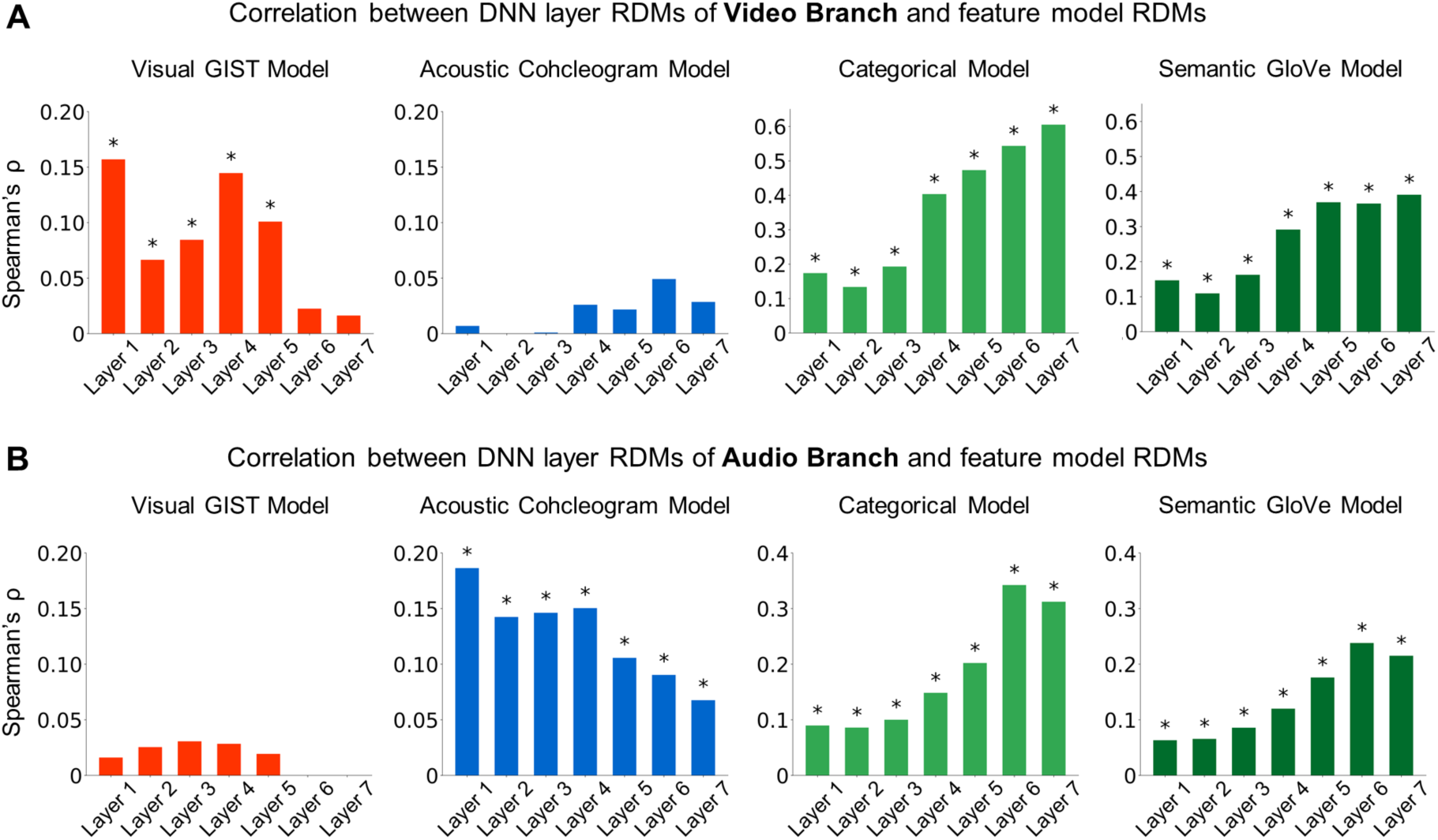
Correlation between layer RDMs of two DNN branches and feature model RDMs. The height of each bar denotes the Spearman’s rank correlation value between two tested model RDMs. The asterisk above the bar represents significant correlations (10,000 permutation test with FDR correction at p<0.01).

## Supplemental Tables

**Table S1:** Descriptions of video stimuli.

| Animals | Objects | Scenes |
| --- | --- | --- |
| cat meowing | alarm ringing | beach |
| cow mooing | blender working | billiard hall |
| cricket chirping | book flipping | bowling hall |
| crown cawing | cellphone vibrating | church |
| dog barking | clock ticking | classroom |
| donkey braying | door opening | firework |
| duck quaking | drum rolling | lake |
| elephant trumpeting | glass clinking | music festival |
| frog croaking | heart monitor beeping | orchestra |
| goose honking | keyboard typing | raining |
| hen clucking | laptop opening | roller coaster |
| horse neighing | money counter working | soccer game stadium |
| lion growling | music box playing | stream |
| mouse squeaking | printer working | street with ambulance running |
| pig oinking | radio tuning | street with car horn |
| pigeon cooing | smoke detector beeping | street with car running |
| sheep bleating | telephone ringing | thunder and lightning |
| rooster crowing | toaster ding | traffic jam |
| sparrow cheeping | toilet flushing | train passing |
| wolf howling | zipper sliding | wind blowing |

